# Joint modeling of zero-inflated longitudinal proportions and time-to-event data with application to a gut microbiome study

**DOI:** 10.1101/2020.12.10.419945

**Authors:** Jiyuan Hu, Chan Wang, Martin J. Blaser, Huilin Li

## Abstract

Recent studies have suggested that the temporal dynamics of the human microbiome may have associations with human health and disease. An increasing number of longitudinal microbiome studies, which record time to disease onset, aim to identify candidate microbes as biomarkers for prognosis. Owing to the ultra-skewness and sparsity of microbiome proportion (relative abundance) data, directly applying traditional statistical methods may result in substantial power loss or spurious inferences. We propose a novel joint modeling framework [JointMM], which is comprised of two sub-models: a longitudinal sub-model called zero-inflated scaled-Beta generalized linear mixed-effects regression to depict the temporal structure of microbial proportions among subjects; and a survival sub-model to characterize the occurrence of an event and its relationship with the longitudinal microbiome proportions. JointMM is specifically designed to handle the zero-inflated and highly skewed longitudinal microbial proportion data and examine whether the temporal pattern of microbial presence and/or the non-zero microbial proportions are associated with differences in the time to an event. The longitudinal sub-model of JointMM also provides the capacity to investigate how the (time-varying) covariates are related to the temporal microbial presence/absence patterns and/or the changing trend in non-zero proportions. Comprehensive simulations and real data analyses are used to assess the statistical efficiency and interpretability of JointMM.

## 1. Introduction

Rapid advances in high-throughput sequencing (HTS) technologies have enabled studies to profile the human microbiome from multiple body sites and demonstrate its link with human health and diseases including obesity, atherosclerosis, and cancer (Turnbaugh et al. 2006, Cho et al. 2012, Kuczynski et al. 2012, Bokulich et al. 2016, Wu et al. 2019, Ganly et al. 2019). Many of these studies use a case-control or cross-sectional study design, in which only one microbiome sample is collected from each subject to examine the association or correlation with the pathological traits of interest. However, recent studies have suggested that human microbiome is temporally dynamic and that longitudinal changes might be disease-associated (Integrative, H. M. P, 2014, Gilbert et al 2016, Paun et al. 2017, Vatanen et al. 2018). Therefore, many studies shift towards prospective longitudinal experimental designs in which the microbiome is sampled at multiple time points (Gilbert et al. 2016, Livanos et al. 2016, Plantinga et al. 2017, Schirmer et al. 2018, Stewart et al. 2018, Schulfer et al. 2019). In these studies, survival (time to event) outcomes are often recorded, allowing investigators to further examine the association of temporal dynamics of microbiome profile with time to event onset and identify signal microbes (Livanos et al. 2016, Plantinga et al. 2017, Stewart et al. 2018).

Unlike other HTS data, microbiome data have specific characteristics that require careful consideration (Xia et al. 2018). 16S rRNA amplicon sequencing and metagenomic shotgun sequencing techniques are the two most widely used HTS approaches to decipher microbiome composition and function. Starting from raw DNA sequencing data, bioinformatics pipelines such as Qiime2 (Bolyen et al. 2019) and MetaPhlAn2 (Truong et al. 2015) can be used for 16S rRNA and metagenomic shotgun sequencing data, respectively, to quantify microbial counts or proportions via aligning reads to reference sequence databases. The microbial counts from 16S rRNA data are not comparable across samples as there is an arbitrary total imposed by the sequencing techniques. Normalizing counts with respect to their total into proportions is a common approach prior to statistical analysis (Gloor et al. 2017). The microbiome proportion data (termed the relative abundance data or proportion data interchangeably herein), either from 16S rRNA amplicon sequencing or metagenomic shotgun sequencing technique(s), is ultra-sparse with many proportions being zero, and the non-zero proportions are highly skewed towards zero, especially for rare taxa (Chen and Li 2016, Peng et al. 2016). Directly applying traditional statistical methods with normal distribution assumptions may lead to underpowered or even spurious results. In the recent years, multiple statistical association analysis methods and software have been proposed to tackle the distribution challenges in microbiome data analyses for both case-control and cross-sectional studies (Li et al. 2015; Koh et al. 2017; Hu et al. 2018). Several mixed-effects models have also been developed for longitudinal or repeatedly measured microbiome data to conduct temporal differential abundance analyses or association studies (Chen and Li 2016; Zhang et al. 2017; Zhang et al. (2) 2017). However, to the best of our knowledge, no models have yet been proposed to jointly model longitudinal microbiome proportion and time-to-event data aiming at examining their associations while taking the special microbiome data structure into account.

Several traditional statistical methods can be modified to analyze longitudinal microbiome proportion and time-to-event data simultaneously, yet all have limitations. One strategy is to treat the longitudinal microbiome proportions as time-dependent covariates and use extended Cox proportional hazards (PH) models to examine the association of event time with time-varying microbial proportions (Fleming and Harrington, 2011). However, the extended Cox model does not consider the randomness of covariates and assumes step-functions for the process of time-dependent covariates, i.e., the microbial proportions remain constant between adjacent time points, which is not realistic. Joint modeling, by treating both longitudinal biomarkers and time to events as outcomes, is another popular and flexible strategy (Faucett and Thomas 1996, Dimitris Rizopoulos (2012)). To investigate the association structure between two different types of outcomes, joint modeling methodologies usually consist of a sub-model for the longitudinal process and a sub-model for the survival process, with linkage via shared latent random effects (Dimitris Rizopoulos (2012)). However, the majority of joint modeling methodologies are developed for normally distributed longitudinal outcomes, while the microbial proportions deviate greatly from the normal distribution assumption, and the deviation does not attenuate after taking the arcsine square root transformation used in several microbiome studies (Web Figure 1 illustrates the distribution fitting for microbiome relative abundance data) (Kostic et al. 2015; Chen and Li, 2016). Moreover, neither of the above methods is able to handle the excessive zeros in the microbiome data analysis. Therefore, a joint modeling method specifically designed to handle the characteristics of longitudinal microbiome proportion data, rigorously model the within-subject correlation of microbiome proportions, and examine its association with clinical or pathological survival outcomes would have substantial potential.

**Figure 1.**
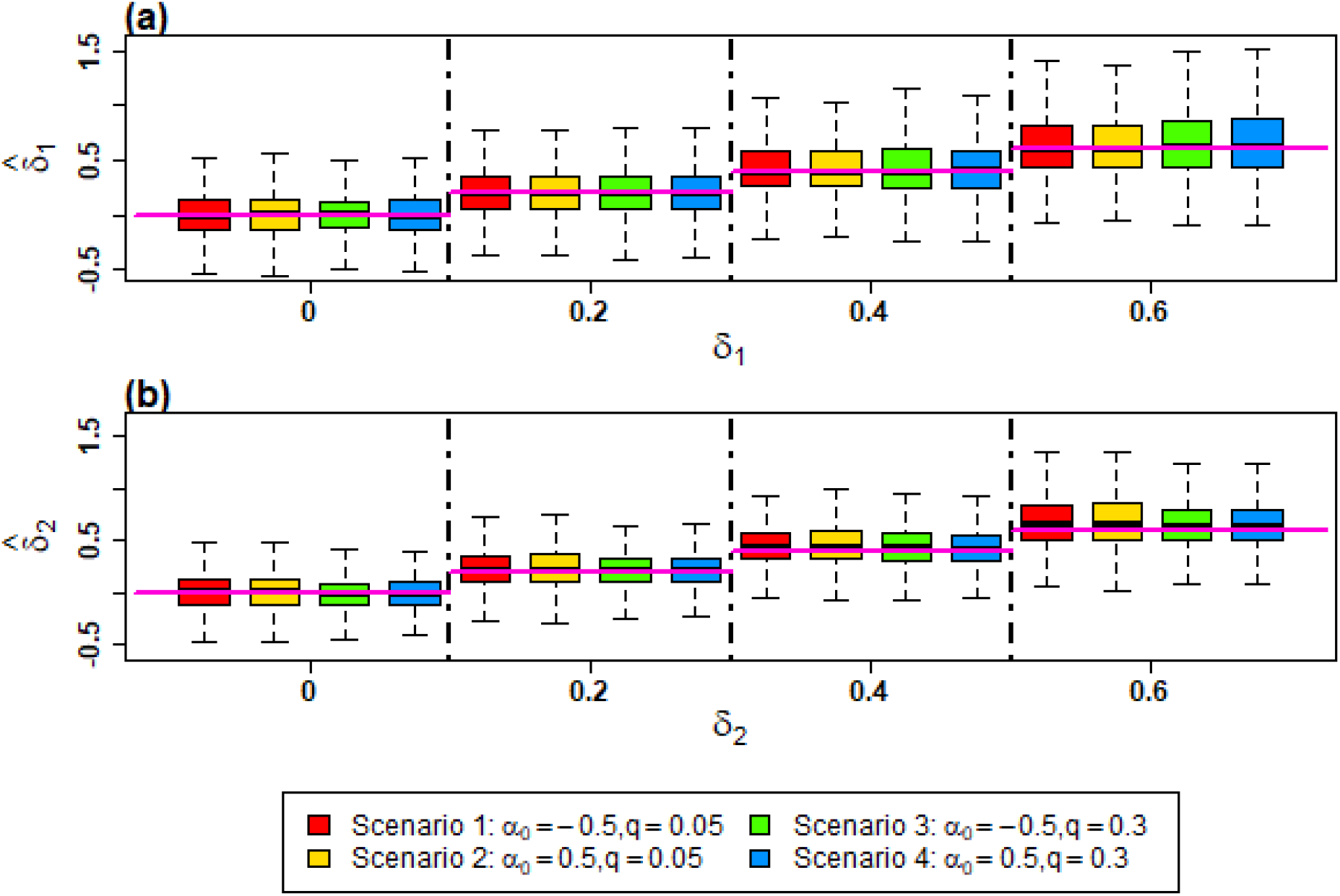
Point estimates for *δ*_1_ and *δ*_2_ in JointMM. Point estimates of 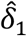in (a) and 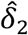 in (b) with *δ*_1_ = *δ*_2_ = 0, 0.2, 0.4, 0.6 respectively under four scenarios for longitudinal microbial proportions and time-to-event association pattern 1. The horizontal purple line indicates the true value of *δ*_1_ or *δ*_2_ under the corresponding scenario.

To address this gap, we propose a joint modeling framework **JointMM** to examine whether temporal microbial presence/absence and/or the non-zero proportion changes are associated with the time to an event, while dealing with the excessive zeros and skewness of microbiome proportion data. JointMM comprises two sub-models: the zero-inflated scaled-Beta (ZIS-Beta) generalized linear mixed-effects model (the longitudinal sub-model) depicts the temporal structure of the microbial proportions among subjects and their associations with confounding factors such as age, gender, and treatment; and the Cox PH model (the survival sub-model) describes the occurrence of an event of interest and its relationship with potential prognostic factors. As one of the key prognostic factors, the longitudinal microbial proportions are linked to the survival outcome via association parameters under the assumption that two sub-models share a latent process (Dimitris Rizopoulos 2012). As a side-product, the longitudinal sub-model of JointMM can also be used alone to investigate the association between (time-varying) covariates and the temporal changes of microbial presence/absence and/or non-zero proportions.

In the next sections, we introduce the notation and the proposed model JointMM (Section 2), and then present the comprehensive simulation results to demonstrate the statistical efficiency and interpretability of JointMM (Section 3). Next, we exhibit how to use JointMM in a real longitudinal gut microbiome study for type 1 diabetes (T1D) (Livanos et al. 2016) to examine the association between the temporal dynamics of gut microbiome proportions and the time toT1D onset and discovered candidate genera whose temporal presence/absence or non-zero proportion changes are T1D-onset-associated(Section 4). Concluding remarks and discussion are in Section 5.

## 2. Methods

### 2.1 Notation and model specification

Suppose we have obtained microbial proportions for all taxa in a longitudinal microbiome study. The proposed model JointMM considers each taxon separately. Let *y*_*ij*_ denote the observed proportion of an arbitrary taxon for subject *i* at time *t*_*ij*_ (*i* = 1, ⋯ , *n*; *j* = 1, ⋯ , *n*_*i*_) and let 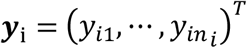 . Note that the observed time points 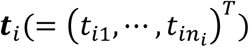 can differ between subjects to accommodate the unbalanced designs. Let 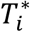 and *C*_*i*_ be the event time and the right-censored time for the *i* th subject, respectively. The observed event time is defined as 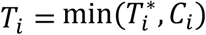 and the event indicator as 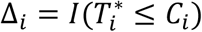, where Δ_*i*_ = 1 if the event is observed and 0 otherwise. In the following, we will sequentially introduce the proposed longitudinal and survival sub-models in JointMM. In accordance with the joint modeling strategy proposed by Henderson *et al*. (2000) and Hickey *et al*. (2018), two sub-models are linked by a multivariate Gaussian latent process.

In the longitudinal sub-model, we first propose a zero-inflated scaled-Beta (ZIS-Beta) mixture distribution to tackle the zero-inflation and skewness of the microbial proportion data. Specifically, the ZIS-Beta distribution for microbial proportion *y*_*ij*_ is given by

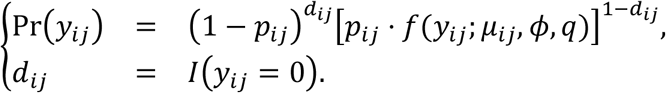

where *p*_*ij*_ represents the probability of the taxon present in the microbiome sample at time *t*_*ij*_ for subject *i* and the probability density function (PDF) of the non-zero part follows the scaled-Beta distribution parameterized as

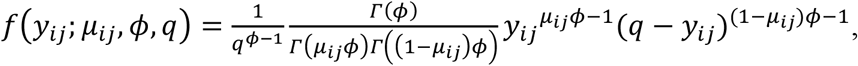

with *μ*_*ij*_(0 < *μ*_*ij*_ < 1) , *ϕ*(*ϕ* > 0) and *q*(0 < *q* ≤ 1) being the mean, dispersion and scale parameters, respectively. Note that the dispersion parameter *ϕ* governs the over-dispersion of proportion data, while the scale parameter *q* ensures the proportion is bounded between 0 and *q*. This is distinct from the Beta distribution which assumes the proportion of any taxon is uniformly bounded between 0 and 1. Considering the sum-to-one constraint in the microbiome data, the proportion of one individual taxon would unlikely range from 0 to 1. Thus, the flexible scaled-Beta distribution assumption suits microbiome data better than the Beta distribution used in other methods (as illustrated in Web Figure 1 B) (Chen and Li, 2016; Peng *et al*., 2016).

Next, we propose a two-part generalized linear mixed-effects model based on the ZIS-Beta distribution to describe the association between covariates and repeated measures of microbial proportions in the following longitudinal sub-model:

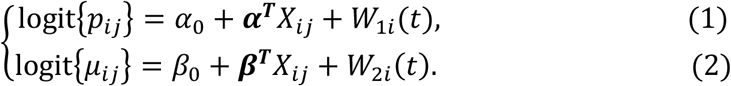

where α_0_ and *β*_0_ are the fixed intercepts indicating the logit transformed baseline presence probability and the non-zero scaled-mean proportion for the studied taxon, respectively; ***α*** and ***β*** are two *p*-vectors of fixed effects of (time-varying) covariates *X*_*ij*_. The Gaussian process ***W***_*i*_(*t*) = (*W*_1*i*_(*t*), *W*_2*i*_(*t*)) ^*T*^ is specified as

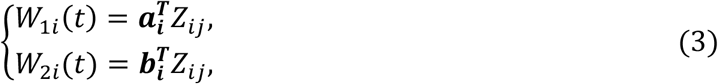

where ***a***_***i***_ and ***b***_***i***_ are the corresponding individual-specific random effects on the *r*-dimensional (time-varying) covariates *Z*_*ij*_, which can be different with *X*_*ij*_. We assume that ***a***_***i***_ and ***b***_***i***_ follow the zero-mean multivariate normal distributions with variance-covariance matrices ***C***_11_ and ***C***_22_, respectively. Since ***a***_***i***_ and ***b***_***i***_ are assumed to be independent for simplicity, the variance-covariance matrix for 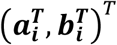 is therefore a block diagonal matrix denoted as 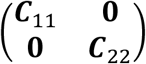. The ZIS-Beta generalized linear mixed effects regression model (1)-(2) depicts the microbial association relationship from two aspects: Equation (1) characterizes the association between (time-varying) covariates and the longitudinal trajectory of microbial presence probability, and Equation (2) links covariates with the temporal trend of non-zero microbial proportions.

For the survival sub-model, we assume that the covariates have multiplicative effects on the hazard for an event and employ the Cox PH model formulated as

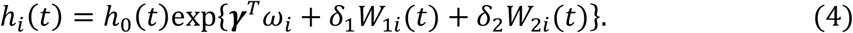

where *h*_0_(.) is the baseline hazard function and ***γ*** corresponds to the coefficients of covariates *ω*_*i*_ which may have association with the survival time. Weibull distribution with shape parameter *k*(*k* > 1) and scale parameter *b*(*b* > 0) is used to model the baseline hazard function *h*_0_(.). The Gaussian processes *W*_1*i*_(*t*) and *W*_2*i*_(*t*) from Equations (1) and (2), respectively, are included in the survival sub-model as the key prognostic factors of interest. With the Gaussian processes shared between the longitudinal sub-model (1)-(2) and the survival sub-model (4), we can investigate the association between the longitudinal microbial proportions and the time-to-event outcome. *δ*_1_ and *δ*_2_ are the corresponding regression coefficients measuring effects of the microbial dynamic pattern in terms of microbial presence probability trajectory and non-zero proportion trajectory, respectively, on the hazard for an event.

In combination, Equations (1), (2) and (4) define the joint model JointMM. Depending on the specification of *Z*_*ij*_ in the Gaussian processes shown in Equation (3), JointMM can investigate random-intercepts (RI) and random-slopes (RS) structures of the longitudinal microbial proportions, respectively, with association to the hazard for an event. With the RI (RS) structure, where within-subject correlation is depicted with random intercepts (random slopes), JointMM can examine the association between the hazard for an event and the subject-specific deviations from the average intercept (slope, i.e., growth rate) of the presence probability/non-zero proportion. It is straightforward to extend JointMM to model random-intercepts and random-slopes (RIRS) structure of the longitudinal microbial proportions theoretically, however, considering the sparsity of microbiome proportion data and limited repeated microbiome measures per subject in many microbiome studies, it is computationally challenging to fit RIRS for JointMM in practical settings. We will illustrate JointMM with RI structure in the following simulation studies and investigate both RI and RS structures separately in our application to assess the gut microbiome for type 1 diabetes.

We also notice that some taxa (especially those dominant ones) have very small fraction of zero microbial proportions, i.e., the assumption of inflation of zeros does not apply. Under such conditions, we only assume the scaled-Beta distribution for the microbial proportions, and remove Equation (1) and the term *δ*_1_*W*_1*i*_(*t*) in Equation (4) accordingly, to build a reduced JointMM. The zero proportions are imputed with the smallest non-zero proportions prior to fit the reduced JointMM. In Section 4, we illustrate the use of full and reduced JointMM in practical settings.

### 2.2 Likelihood

We postulate the widely-adopted conditional independence assumption in the joint modeling research field to determine the likelihood function based on the observed outcome{(*T*_*i*_, *Δ*_*i*_, ***y***_*i*_), *i* = 1, ⋯ , *n*} (Dimitris Rizopoulos 2012). To be specific, the longitudinal and time-to-event data generating processes are conditionally independent on the latent Gaussian process {***W****_i_*(*t*), *i* = 1, ⋯ , *n*} and the observed covariates in the longitudinal and survival sub-models. The parameters of JointMM are denoted by 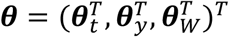, with 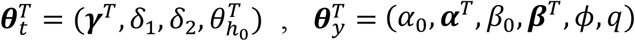 and 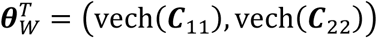, corresponding to the parameters in the survival sub-model, longitudinal sub-model, and the variance-covariance matrix of the latent process, respectively. 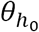 denotes the parameters in the baseline hazard function *h*_0_(.), and vecH(.) denotes the half-vectorization of a matrix. Then the likelihood function can be written as

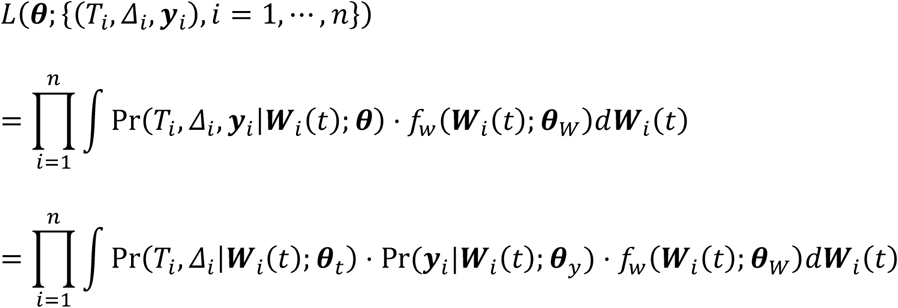

where *f*_*w*_(.; ***θ***_*W*_) is the PDF of the random process ***W***_*i*_(*t*) . Therefore, we can partition the likelihood into two parts, the longitudinal part and the survival part, which can be obtained through the likelihood of their corresponding sub-models (refer to Web Appendix A for the detailed derivation of the likelihood). Gaussian Quadrature rules are used to computationally approximate the integrals involved in the likelihood of the joint model. In practice, the scale parameter *q* is estimated with the largest observed proportion value plus an ignorable small value *ε*. For other parameters, Quasi-Newton constrained optimization algorithm “L-BFGS-B” is employed to seek for the maximum likelihood estimates (MLEs) and the corresponding estimated information matrices (Byrd et al. 1995).

### 2.3 Hypothesis tests

Based on the likelihood in Section 2.2, we use the Wald test to evaluate the following statistical hypotheses. Our primary interest is whether the temporal heterogeneity in presence probability or non-zero proportions among subjects are associated with the time to an event, i.e.,

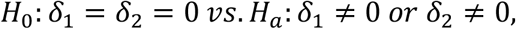

which is regarded as the global null test. JointMM can separately examine whether the temporal heterogeneity of the presence probability of the taxon would affect the event time by

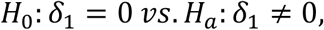

and whether the temporal heterogeneity of the taxon’s non-zero proportion is associated with the survival time by

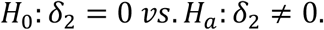

We denote testing of the above hypotheses with JointMM as JointMM-global, JointMM-logit and JointMM-proportion, respectively.

Within the longitudinal sub-model (1)-(2), we can also test if the *k*th covariate (*k* = 1, ⋯ , *p*) is associated with the longitudinal microbial presence trajectory using the null hypothesis *H*_0_:*α*_*k*_ = 0, the microbial proportions trajectories via testing the null hypothesis of *H*_0_:*β*_*k*_ = 0, or either of the two via testing *H*_0_:*α*_*k*_ = *β*_*k*_ = 0.

## 3. Simulation studies

We conducted extensive simulation studies to evaluate the performance of JointMM in both the joint model inference and the microbial longitudinal inference. In the joint model inference for the association examination between longitudinal microbiome and time-to-event outcome, we compared JointMM with two competing methods: (1) the traditional joint modeling approach which is built upon normal distribution assumption (Dimitris Rizopoulos (2012)), and (2) the extended Cox model where longitudinal microbiome proportions are included as time-varying covariates (Fleming and Harrington, 2011). In the microbial longitudinal inference for the association between longitudinal microbial proportions and time-varying covariates, we compared the longitudinal sub model of JointMM with a two-part zero-inflated Beta random effects model (ZIBR) developed by Chen and Li (2016), and the classic linear mixed effects (LME) regression model, respectively. Microbial proportion data were arcsine square root transformed in the models requiring normal distribution assumptions, as suggested in La Rosa *et al*. (2014) and Kostic, et al. (2015).

### 3.1 Joint model inference for longitudinal microbial proportions and time-to-event outcome

#### 3.1.1 Simulation design

We generated longitudinal microbial proportions and time-to-event data with the following RI structure model.

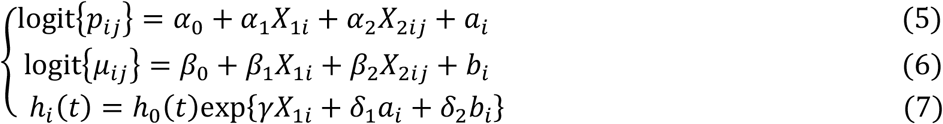

Two covariates were considered in this simulation setting. X_1*i*_ denotes a binary covariate such as gender, where 50% of X_1*i*_s were randomly assigned to 0 and the remaining were assigned to 1. *X*_2*ij*_ = *j* is a time-varying covariate to describe the time pattern, where *j* = 0, 1, ⋯ ,4 for five time points (*i* = 1, ⋯ , *n*). The intercept *α*_0_ and the scale parameter *q* of ZIS-Beta distribution control the fraction of zeros and the upper bound of the microbial proportion data, respectively, based on which we considered four scenarios of generating microbiome proportion data to evaluate comprehensively how the distribution of microbial proportions affect the association tests between two outcomes.

▪ **Scenario 1**: α_0_ = −0.5, *q* = 0.05 which corresponds to data with high zero fractions (∼53%) and low non-zero microbial proportions;
▪ **Scenario 2**: α_0_ = 0.5, *q* = 0.05 which corresponds to data with low zero fractions (∼34%) and low non-zero microbial proportions;
▪ **Scenario 3**: α_0_ = −0.5, *q* = 0.30 which corresponds to data with high zero fractions (∼53%) and high non-zero microbial proportions;
▪ **Scenario 4**: α_0_ = 0.5, *q* = 0.30 which corresponds to data with low zero fractions (∼34%) and high non-zero microbial proportions.

Scenario 1 represents the typical distributions of rare taxa, where taxa are present in a few samples with low relative abundances. Scenario 4 deals with the opposite case involving common taxa. Scenario 2 represents those taxa presenting in most samples but with very low relative abundances, whereas Scenario 3 represents taxa which are present only in a few samples at high relative abundances. We set the other regression coefficients in the longitudinal sub-model as(*α*_1_, *α*_2_)^*T*^ = (*β*_1_, *β*_2_)^*T*^ = (0.5,0.05)^*T*^ and *β*_0_ = −3.0. The dispersion parameter of ZIS-Beta distribution was generated as *ϕ* ∼ Uniform(2,10). The shared random effects (*a*_*i*_, *b*_*i*_) were assumed independent with standard deviation of the normal distribution with *σ*_1_ = *σ*_2_ = 1.

For the time-to-event random process, we included the covariate X_1*i*_ with regression coefficient *γ* = 0.5 and considered three patterns of longitudinal microbiome and time-to-event associations:

1. Both the temporal heterogeneity of presence probability and the non-zero proportions are associated with the event time, with effect size *δ*_1_ = *δ*_2_ = *δ* = 0, 0.2, 0.4 or 0.6 (*δ* = 0 is used to evaluate the type I error rates);
2. Only the temporal heterogeneity in microbial presence probability is associated with the event time with effect size *δ*_1_ = 0.2, 0.4 or 0.6, *δ*_2_ = 0;
3. Only the temporal heterogeneity in the microbial taxon’s non-zero proportions is associated with the event time with effect size *δ*_1_ = 0, *δ*_2_ = 0.2, 0.4 or 0.6.

The baseline hazard *h*_0_(*t*) follows a Weibull distribution with shape and scale parameters *k* = 1.05, and *b* = 0.05 respectively. Independent censoring is set as *C* = Uniform(0, 6), representing ∼40% of censoring for event time.

Longitudinal microbiome and time-to-event data were simulated for *n* = 100 subjects and we conducted 5,000 independent repetitions to evaluate the estimation of parameters, and the type I error rates and power for association tests of JointMM-global, JointMM-logit and JointMM-proportion, respectively, under three longitudinal microbiome and time-to-event association patterns at significance level of 0.05. Our simulation setup is designed based on the joint modeling framework for analyzing medical cost data proposed by Lei Liu (2009) and the mixed-effects model ZIBR developed by Chen and Li (2016) for longitudinal microbiome proportion data.

#### 3.1.2 Competing methods

We compared JointMM with the traditional joint modeling method JM for analyzing normally distributed longitudinal data and the extended Cox regression model (Dimitris Rizopoulos 2012; Fleming and Harrington, 2011). The arcsine square root transformed longitudinal microbial proportions are denoted for subject *i* by 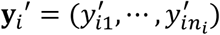. The joint model JM with RI structure for simulation data in Section 3.1.1 is given by

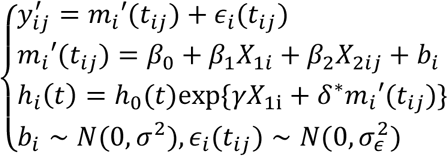

where the random effects *b*_*i*_ and error term *∈*_*i*_(*t*_*ij*_) are independent. We used the function **jointModel()** from R package **JM** with parameter **method = “weibull-PH-GH”** to fit the model and evaluate the likelihood ratio test for *H*_0_:*δ*^∗^ = 0 (Dimitris Rizopoulos 2012). We also fitted data with the extended Cox model with R package **survival** to examine the association of event time with time-varying microbial proportions with Wald test by treating longitudinal microbial proportions as a time-varying covariate. The above methods are denoted by JM and ExtendCox, respectively, hereafter for simplicity.

#### 3.1.3 Simulation results of joint inference

We first evaluate the estimation of two association parameters of primary interest in JointMM, i.e., *δ*_1_ and *δ*_2_. Figure 1 displays the corresponding point estimates for four Scenarios of microbial proportion distribution under the first longitudinal microbiome and time-to-event association pattern. The estimation bias of 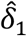 and 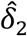 is negligible under most of the conditions, except that very minor upward bias of 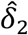 is depicted when *δ*_1_ = *δ*_2_ = 0.6 under Scenario 1. This is mainly due to the ultra-sparsity and low non-zero relative abundances of the microbial proportion data under Scenario 1 as there are over half of microbial proportions being zero and the maximum relative abundance is less than *q=*0.05 (the upper bound). The point estimates of *δ*_1_ and *δ*_2_ with the other two association patterns are similar and thus not shown here.

Next, we compare the testing results of JointMM-global, JM and ExtendCox regarding the global null hypothesis for three association patterns described in Section 3.1.1. We also evaluate the association testing performance of JointMM-logit and JointMM-proportion with reference to the longitudinal trajectory of presence probability and non-zero proportions, respectively. Figure 2 shows the type I error rates (when *δ*_1_ = *δ*_2_ = 0) and power (when *δ*_1_ = *δ*_2_ = 0.2, 0.4, 0.6) under four Scenarios in which both the longitudinal presence probability and non-zero proportions of taxa are event-time associated (association pattern 1). The type I error rates of five testing methods are all well-controlled around the nominal level of 0.05. JointMM-global achieves uniformly the highest power on testing the global null under four scenarios compared with JM and ExtendCox. JM exhibits lowest power in Scenario 1 and 2, where the microbial proportions are highly-skewed towards 0 with the maximum relative abundance <0.05 and are far from normal even with arcsine square root transformation. The statistical power of JM is higher in Scenarios 3 and 4 which model more abundant microbial proportions and lower zero fractions than in Scenarios 1 and 2. ExtendCox is robust across four Scenarios, showing relatively lower or similar power with JM. The robustness of ExtendCox is due to its model assumption that the longitudinal microbial proportions are included as time-varying covariates, hence the distribution changes of the proportions has less impact on the testing result of ExtendCox compared with JM. JointMM-logit achieves better performance regarding testing of the presence/absence pattern under Scenario 1 and 3 where the fraction of zero microbial proportions is higher and sufficient microbial proportion data can be used to fit the logit model in Equation (1) compared with its results under Scenario 2 and 4. The power of JointMM-proportion is similar under Scenario 1 and 3, and under Scenario 2 and 4. Note that the main differences between Scenario 1 and 3 (or between Scenario 2 and 4) are the distribution range of non-zero proportions. JointMM-proportion is insensitive to the microbial proportions, which indicates the usefulness of including the scale parameter in modeling the microbial proportion distribution.

**Figure 2.**
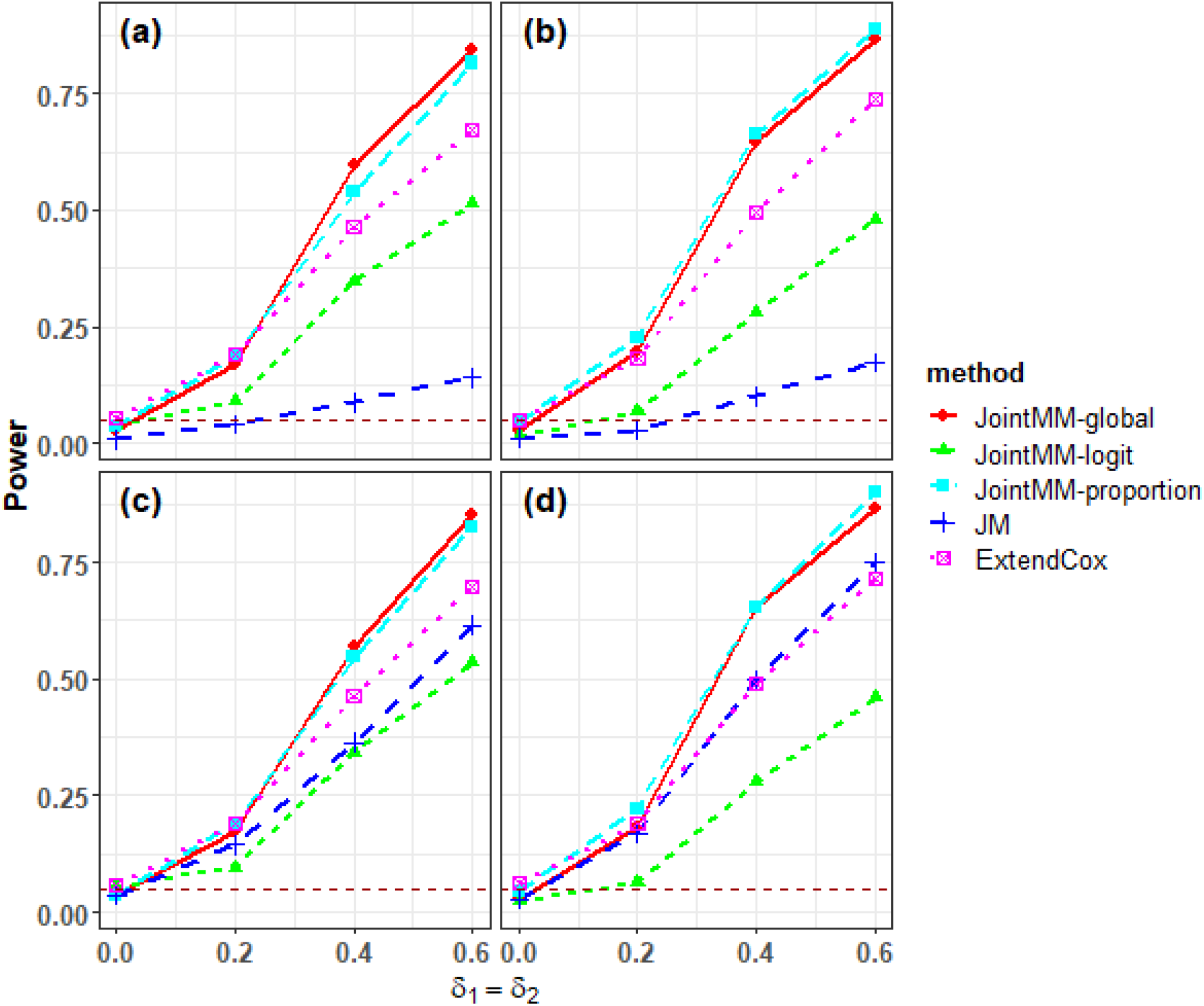
Longitudinal microbial proportions and time-to-event association pattern 1. Shown are the type I error rates (*δ*_1_ = *δ*_2_ = 0) and statistical power (*δ*_1_ = *δ*_2_ = 0.2,0.4,0.6) for testing the global null hypothesis with JointMM-global, JM and ExtendCox, respectively, and for testing the association of the longitudinal presence probability and non-zero proportions with the event time with JointMM-logit and JointMM-proportion, respectively. Results under Scenarios 1-4 are shown in panels (a)-(d), respectively (significance level = 0.05).

The association testing results when only the temporal microbiome presence probability (association pattern 2) or the non-zero proportions (association pattern 3) is associated with time to an event are depicted in Figures 3 and 4, respectively. Under association pattern 2 (δ_2_ = 0, *δ*_1_ = 0.2, 0.4,0.6), JointMM-global achieves the highest power with reference to testing the global null hypothesis for four Scenarios. In comparison, JM almost fails in detecting the association since the microbial proportions JM used for statistical inference (the non-zero observations) are in fact not associated with the survival outcome. ExtendCox also has limited statistical power. The JointMM-proportion in Figure 3 shows well-controlled type I error rates ∼0.05 for all conditions since δ_2_ = 0 under all four scenarios of association pattern 2. Under association pattern 3 (δ_1_ = 0, *δ*_2_ = 0.2, 0.4,0.6) (Figure 4), JointMM-global continues to have the highest power as it models the longitudinal microbial proportion data from two aspects, i.e., presence/absence and non-zero proportions. JointMM-logit illustrates a statistical valid performance that the empirical type I error rates are around nominal level for all conditions.

**Figure 3.**
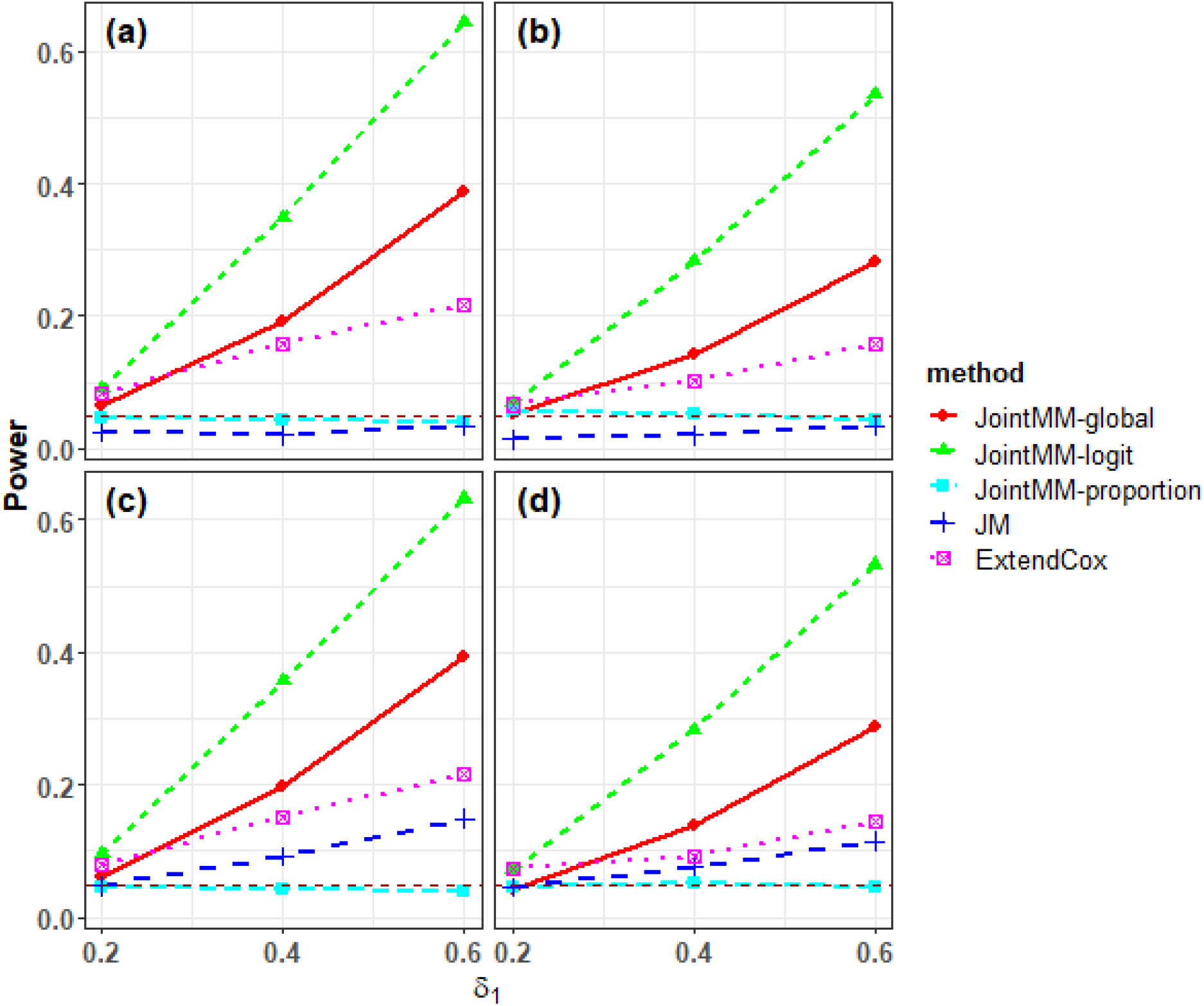
Longitudinal microbial proportions and time-to-event association pattern 2. Shown are the statistical power (*δ*_1_ = 0.2, 0.4, 0.6, *δ*_2_ = 0) for testing the global null hypothesis with JointMM-global, JM and ExtendCox, respectively, and for testing the effect of the longitudinal presence probability and (non-zero) proportions on the event time with JointMM-logit and JointMM-proportion respectively. Results under Scenarios 1-4 are shown in panel (a)-(d), respectively (significance level = 0.05).

**Figure 4.**
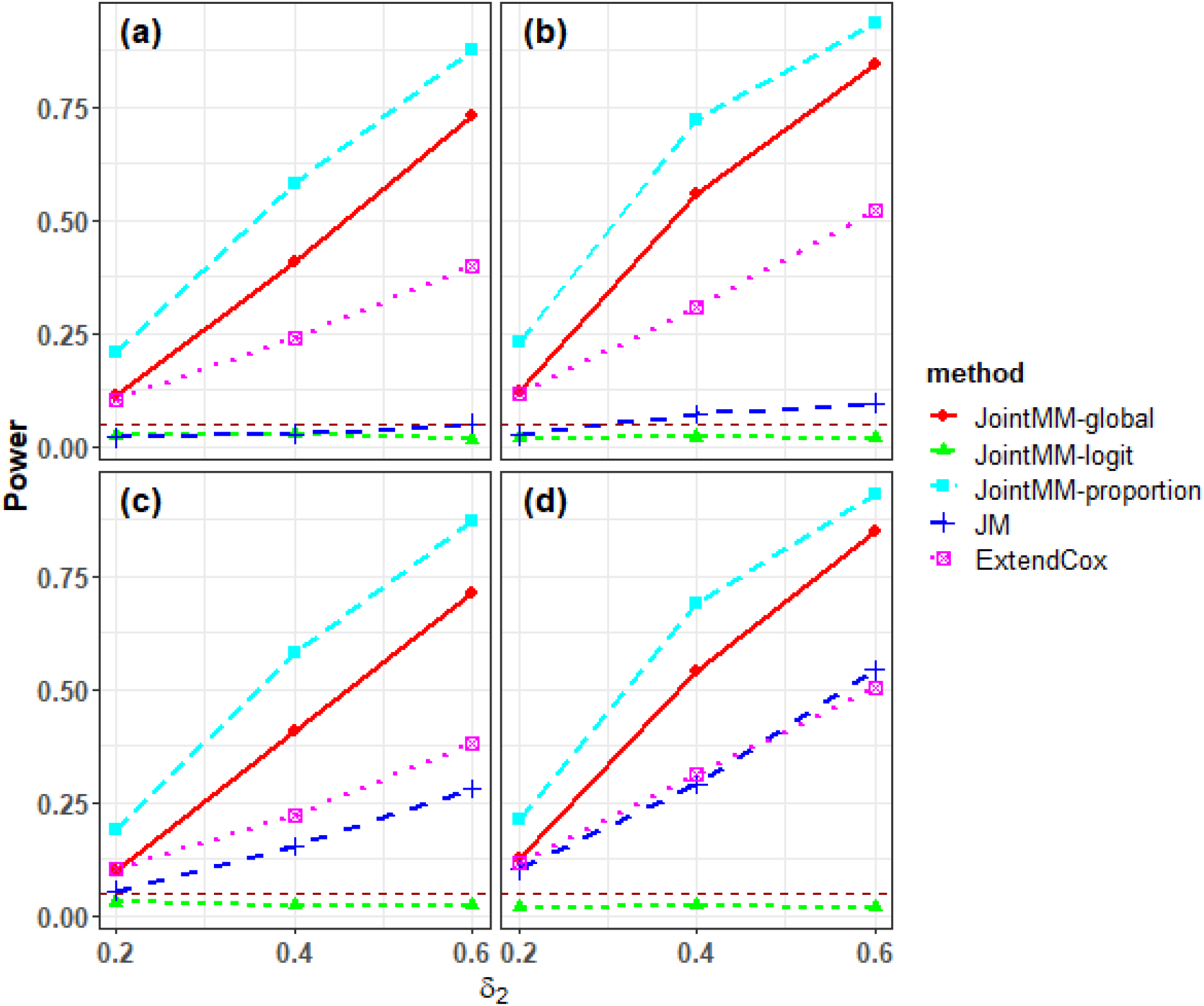
Longitudinal microbial proportions and time-to-event association pattern 3. Shown are the statistical power (*δ*_2_ = 0.2, 0.4, 0.6, *δ*_1_ =0) for testing of the global null hypothesis with JointMM-global, JM and ExtendCox respectively, and for testing of the effect of the longitudinal presence probability and (non-zero) proportions on the event time with JointMM-logit and JointMM-proportion respectively. Results under Scenarios 1-4 are shown in panels (a)-(d) respectively (significance level = 0.05).

In conclusion, JointMM-global achieves uniformly greater power compared with JM and ExtendCox. Modification of traditional methods, such as applying JM to arcsine square root transformed microbiome proportion data, or treating the abundance data as covariates as in ExtendCox would lead to significant power loss. In addition, the testing of JointMM-logit and JointMM-proportion provide further delineation of the association patterns.

### 3.2 Inference of the longitudinal sub-model

We further evaluated the statistical inferences of the longitudinal sub-model of JointMM, compared with the two-part mixed-effects model ZIBR developed by Chen and Li (2016) and the classic linear mixed effects (LME) regression model to assess the statistical validity. ZIBR also incorporates a two-part model to infer whether the covariates are associated with the temporal trajectories of microbial presence probability and/or proportions. Microbial proportions are arcsine square root transformed prior to fitting the LME model.

Longitudinal microbiome proportion data were generated from Equation (5)-(6) based on the parameters established, as follows. Only the binary covariate X_1*i*_ was included for simplicity and three microbial association scenarios are considered.

▪ **Scenario 5**: α_1_ = *β*_1_ = 0.5 where covariate X_1*i*_ affects both the temporal microbial presence probability and non-zero proportions;
▪ **Scenario 6**: α_1_ = 0.5, *β*_1_ = 0 where covariate X_1*i*_ only affects the longitudinal microbial presence probability;
▪ **Scenario 7**: α_1_ = 0, *β*_1_ = 0.5 where covariate X_1*i*_ only affects the longitudinal microbial non-zero proportions trajectory.

The intercept terms were set as(*α*_0_, *β*_0_)^*T*^ = (−0.5, −3)^*T*^, scale parameter *q* = 0.3 to illustrate common taxa and all the other parameters were generated as illustrated in Section 3.1.1. Statistical power of association tests for the global null hypothesis of H_0_: *α*_1_ = *β*_1_ = 0 with JointMM, ZIBR and LME, and association tests of the null hypotheses H_0_:*α*_1_ = 0 (microbial presence) and H_0_:*β*_1_ = 0 (microbial non-zero proportions) with JointMM and ZIBR were evaluated at significance level 0.05.

Web Figure 2 illustrates the statistical power of three competing methods. Under Scenario 5, where the covariate affects both the longitudinal microbial presence probability and non-zero proportions with equal association strength (regression coefficients), LME has relatively higher power in testing for the global null hypothesis compared with JointMM and ZIBR. However, the power of LME is evidently lower than that of JointMM and ZIBR when the covariate only affects the microbial presence (Scenario 6) or non-zero proportions (Scenario 7), which is more commonly observed in real data conditions. The longitudinal sub-model of JointMM achieves greater power than ZIBR in testing for all three hypotheses, which may due to the introduction of a scale parameter in JointMM that better deals with the skewness of microbial proportion data. Both JointMM and ZIBR successfully control the empirical false positive rates under the nominal lever of 0.05 for testing the association of microbial presence (microbial non-zero proportions) under Scenario 6 (Scenario 7).

## 4. Application to a gut microbiome study in T1D

We applied the proposed joint modeling framework JointMM to a longitudinal T1D study with non-obese diabetic mice. The study investigated how the early-life antibiotic usage perturbed the gut microbiome and their relationship with the onset of T1D (Livanos et al. 2016). In this study, new born mice were randomized tocontinuous low-dose (sub-therapeutic) antibiotics (STAT), pulsed therapeutic antibiotic dosing (PAT), or no antibiotics (control) group, and were tested for diabetes weekly from week 10 until week 31 when the follow-up ended. Censoring of the onset of T1D (right censoring) occurred if mice were sacrificed to obtain other biological samples prior to T1D onset or were T1D-free when the study ended. Their fecal samples from week 3, 6, 10 and 13 were collected and 16S rRNA amplicon sequencing was conducted for the microbial DNAs, as detailed in Caporaso et al. (2012) and Livanos et al. (2016). In total, 352 microbiome samples were obtained from 98 mice, resulting in an average of 3.6 repeated measures of microbiome composition per subject. 106 genera were observed originally using the QIIME pipeline (Caporaso et al. 2010) and then were removed if the genera i) have unidentified lineage, ii) were present in less than 10% of samples, or iii) have average microbial proportions < 0.01%. After filtering, the proportion data relative to 19 genera were retained for our analyses.

Since Livanos et al. (2016) has demonstrated that STAT and control samples in both sexes at all timepoints substantially overlap in community structure measured by β-diversity, and the T1D onsets in these two groups have no significantly different pattern (Figures 4 and 1 in Livanos et al.; 2016), we therefore pooled the microbiome samples from STAT and control groups to investigate how the temporal presence probability and/or non-zero proportions of bacterial genera in control/STAT mice are associated with time-to-T1D onset. Both RI and RS random effects structures were investigated respectively with JointMM. JointMM and competing methods were fitted for the microbial proportion data of each genus and time-to-T1D onset and then the Benjamini-Hochberg procedure was applied to the raw p-values to control for the false discovery rate (FDR = 0.05) (Benjamini and Hochberg, 1995). Specifically, we fitted JointMM and JM with antibiotic treatment, sex and sampling time (in weeks) as covariates of the longitudinal sub-model; and antibiotic treatment and sex as covariates in the survival sub-model. The random effects structure of RI were investigated first for the joint inference. The reduced model of JointMM was fitted to genera where over 90% the observed microbial proportions are non-zero. Treatment, sex, and longitudinal proportions (arcsine square root transformed) were included as (time-varying) covariates in ExtendCox for comparison. The longitudinal microbiome-T1D onset association testing for the global null with JointMM-global, JM and ExtendCox is shown in Table 1. JointMM identified five genera including *Turicibacter, Enterococcus, Eubacterium, Anaerostipes* and *Bifidobacterium* whose temporal microbial presence and/or non-zero proportions are associated with the time to T1D onset (FDR = 0.05). In comparison, JM and ExtendCox did not detect any genus whose temporal proportions were significantly associated with T1D onset. Noticeably, JM failed to converge when fitting the transformed proportion data of *Enterococcus*. Testing results of the global null in the joint inference demonstrate marked performance of JointMM which is consistent with our simulation studies. In contrast, the traditional methods JM and ExtendCox, have limited power for inference of longitudinal microbiome-survival association detection. In addition (Table 1), JointMM-logit identified three genera (*Turicibacter, Enterococcus, Eubacterium*) whose temporal presence probability trajectory was associated with T1D onset, and JointMM-proportions identified two other genera (*Anaerostipes, Bifidobacterium*) whose longitudinal microbial proportion changes were associated with T1D onset. We then fit JointMM with the same covariates and investigated the random effects of RS structure to detect candidate gut bacteria whose growth rate is associated with T1D onset. Web Table 1 shows the testing results for JointMM-global, JointMM-logit, and JointMM-proportion, respectively, and we found substantial overlap of candidate bacteria that their longitudinal microbial presence and/or non-zero proportions were associated with T1D onset under RI and RS structures.

**Table 1.**
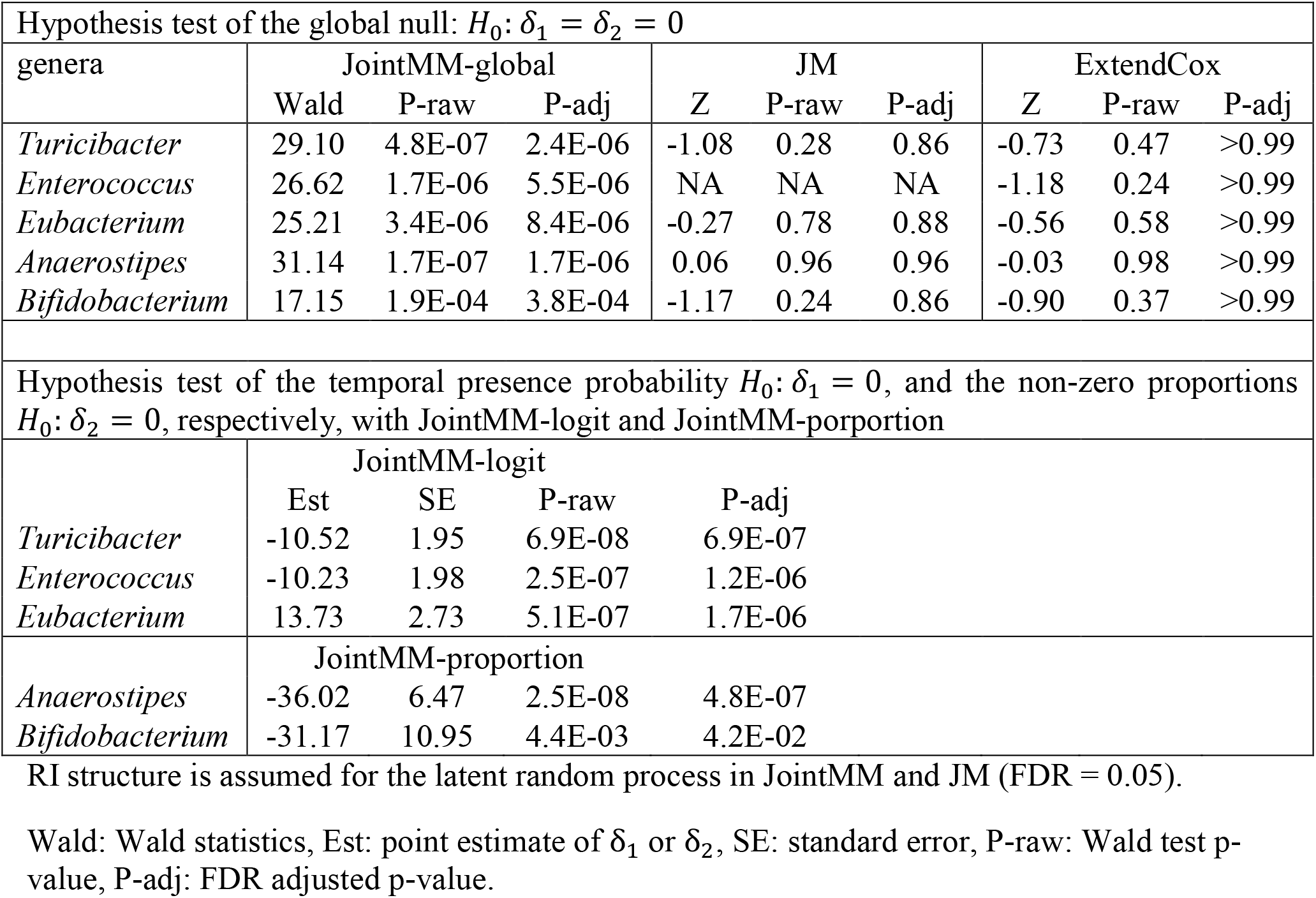
Joint inference of the association between longitudinal microbiome and time to T1D onset data (Livanos et al. 2016) with JointMM-global, JM, and ExtendCox, respectively.

We also investigated the inference results of the longitudinal sub-model of JointMM and compared with ZIBR and LME in analyzing longitudinal microbiome data. Web Table 2 illustrates the genera which were detected by the longitudinal sub-model of JointMM, ZIBR, or LME to have temporal differential presence/absence and/or non-zero proportions between STAT and Control treatment group, after adjusting for gender and time effects. JointMM identified three and two genera whose temporal presence probability and non-zero proportions, respectively, differ between the two treatment groups. The association patterns (temporal differential presence or proportions) identified by ZIBR is consistent with that of JointMM, which further demonstrates the validity of JointMM. LME detected four temporal differential abundance genera, however it cannot distinguish the association patterns which can be distinguished by JointMM and ZIBR. Web Figure 3 shows the observed and predicted presence probability (logit transformed) and non-zero proportions (logit transformed) over time for five significant genera. In the STAT group, *Anaerostipes, Roseburia* and *Turicibacter* have lower probability of presence, and *Lactobacillus* is less abundant. This may indicate that these genera are sensitive to the antibiotic exposure (STAT) during the early life of the host. Among them, four genera have been shown to become more prevalent (*Anaerostipes, Roseburia*) or are more abundant (*Lactobacillus, Bifidobacterium*) over time, showing a trend of microbial recovery from the antibiotic disturbance.

## 5. Discussion

In this paper, we propose a novel joint modeling framework JointMM specifically for analyzing longitudinal microbiome proportions and time-to-event data. JointMM is capable of investigating the effects of longitudinal microbial presence and non-zero proportion patterns respectively on the time-to-event outcome, and can serve as a perfect tool to detect microbes as candidate biomarkers for disease prognosis, which will help to elucidate the role of the human microbiome in human health. To be specific, JointMM can identify microbial taxa whose temporal presence probability and/or non-zero proportion are associated with the hazard for the disease via introducing the latent random effects shared between the longitudinal microbial proportion process and the time-to-event process. We propose a ZIS-Beta mixture distribution along with the ZIS-Beta generalized linear mixed effects model to tackle the zero inflation and skewness of longitudinal microbial proportion data. JointMM is flexible to use that permits inferences about associations between longitudinal microbiome proportions and (time-varying) covariates, by using the corresponding longitudinal sub-model of JointMM. Comprehensive simulation studies and real data application show the statistical efficiency of JointMM.

In the joint modeling arena, Expectation-Maximization (EM) algorithm is often used in maximizing the log-likelihood function with respect to the parameters since some of the parameters have closed-form updates in the M-step. For instance, Rizopoulos, D. (2012) reported the closed-form expressions for the measurement error variance in the longitudinal sub-model and the random effects covariance matrix for JM under normal distribution assumptions in the M-step. In JointMM, however, there is no closed-form expression in the M-step of EM algorithm due to the mixture distribution and the generalized linear mixed effects regression model introduced herein to tackle the zero inflation and skewness of microbiome proportion data. Therefore we use Quasi-Newton constrained optimization algorithm instead for the optimization of the log-likelihood function. To accelerate the convergence of optimization, we first fit two sub-models (random processes are not included in the survival model) to the longitudinal microbiome data and survival data separately, and adopt the corresponding parameter estimates as the initial parameter value for optimizing the likelihood of the joint model.

Currently we have focused on developing statistical models to better characterize microbiome proportion data and proposing association tests to more powerfully identify candidate taxa whose longitudinal proportions are associated with the time-to-event outcome. Microbiome research can surely be advanced by the availability of individualized predictions of subject prognosis based on their longitudinal microbiome profiles. While association analysis emphasizes the modeling of population-level trajectories, personalized prediction requires higher flexibility of the model that can account for the nonlinear patterns in each individual’s longitudinal trajectory. As a future research area, we will aim to extend JointMM to predict the time to an event for each individual based on the longitudinal microbiome profile of candidate microbes and additional information about covariates. Non-linear modeling techniques such as piecewise linear trajectory for each individual could be incorporated to more flexibly capture each subject’s coevolution of longitudinal and time-to-event processes (Barrett and Su, 2016).

JointMM performs longitudinal microbiome and time-to-event association screening in two steps: first modeling the longitudinal proportions of each microbial taxon and the time-to-event outcome and then controlling the false discovery rate for all taxa among the profiled microbiome. Another potential research direction is to incorporate all taxa of the microbiome profile into a single model and jointly investigate the effects of the temporal abundances of all microbes (multiple longitudinal outcomes) on the time to an event. Nonetheless, multivariate joint modeling (MVJM) methods face severe computational challenges due to the requirement for numerical integration of the random effects (Dimitris Rizopoulos 2012; Hickey et al. 2016). Computational limitations would preclude large numbers of random effects which increase exponentially with the number of taxa. Regularization approaches are therefore of potential for MVJM methods, considering the tens to hundreds of microbial taxa confronted in microbiome data analysis.

In conclusion, we provide a useful joint modeling method for detecting the association between longitudinal microbiome and the time-to-event outcome. The proposed joint modeling framework has been implemented in R package JointMM and is publicly available at https://sites.google.com/site/huilinli09/software and https://github.com/JiyuanHu/JointMM.

## 6. Acknowledgements

This work was supported in part by National Institutes of Health grants R01DK090989, R01DK110014 and U01AI22285, and the Zlinkoff and C&D Funds.

## Supporting Information

Additional supporting information may be found online in the Supporting Information section at the end of the article.

